# Modeling natural HIV-1 infection and immunopathogenesis in a lymphoid organ-chip

**DOI:** 10.64898/2026.11.27.769475

**Authors:** Abdur Rashid, Cinthya Zelaya Aparicio, Daniel Ramos, Matthew L. Anderson, Brittney Short, Charles Rivera, Leigh Anne Eller, Nelson L. Michael, Merlin L. Robb, Gautam Mahajan, Hongshuo Song

## Abstract

Secondary lymphoid organs are the principal sites of HIV-1 replication, immune dysfunction and reservoir persistence, yet physiologically relevant human models of this microenvironment remain limited. Here we show that a microfluidic lymphoid organ-chip supports productive HIV-1 replication without exogenous T cell activation or cytokine supplementation. All six HIV-1 strains tested, including three CXCR4-tropic (X4) and three CCR5-tropic (R5) viruses, replicated productively in the chip, whereas matched static three-dimensional cultures supported only three. Infection recapitulated key features of HIV-1 immunopathogenesis, including CD4 T cell depletion, a decreased CD4/CD8 ratio, loss of T follicular helper cells, and altered T cell turnover and activation. The chip also reproduced the greater pathogenicity of X4 over R5 HIV-1, concordant with observations in natural HIV-1 infection, and revealed previously unidentified divergent impacts of X4 and R5 strains on CD4 T cell proliferation and activation. Antiretroviral therapy rapidly suppressed HIV-1 replication, reduced reservoir size, and reversed infection-induced T cell phenotypic changes. The lymphoid organ-chip thus provides an immunocompetent, human relevant model in which HIV-1 infection and host immunity can be studied simultaneously.

## Introduction

More than four decades into the HIV-1 pandemic, an effective vaccine and a functional cure remain unavailable. Although research over the past decades has substantially advanced our understanding of HIV-1 virology and immunology, fundamental questions persist regarding HIV-1 transmissibility, immunopathogenesis, reservoir persistence and the mechanisms underlying incomplete immune reconstitution during antiretroviral therapy (ART). Addressing these questions will be essential to inform interventions towards effective HIV-1 prevention and functional cure.

Research in the HIV-1 field has long been constrained by the absence of a physiologically relevant human system that mimics the microenvironment of secondary lymphoid organs (SLOs), the principal sites of HIV-1 replication, immune dysfunction and reservoir persistence. Recreating the SLO microenvironment *ex vivo* has proved challenging, and each available human model has limitations. Two-dimensional (2D) cultures of peripheral blood mononuclear cells (PBMCs) or purified CD4 T cells support high-level HIV-1 replication but require exogenous activation. An important limitation is that artificial T cell activation causes the loss of T cell subset heterogeneity, which is critical for understanding HIV-1 pathogenesis, transmissibility and reservoir composition^1,2^. *Ex vivo* lymphoid tissue models overcome the requirement for exogenous activation. Human lymphoid histocultures (HLH)^3^ and lymphoid aggregate cultures (HLAC)^4^ both support productive HIV-1 replication without exogenous stimulation, and both have provided important insights into HIV-1 pathogenicity^5–8^. A limitation shared by both models is that the cultures typically last no longer than two weeks^4,9^. Tonsil organoids, first developed by Wagar et al.^10^, offer improved longevity and reconstitute functional germinal center-like structures. To date, however, evidence that tonsil organoids support productive replication of diverse primary HIV-1 strains without exogenous T cell stimulation remains minimal: among studies exposing tonsil organoids to replication-competent HIV-1, only two have tracked viral replication kinetics over time in a manner indicating successive rounds of replication without exogenous activation^11,12^. Critically, HLH, HLAC and organoid cultures are all maintained under static three-dimensional (3D) conditions and therefore cannot recapitulate the dynamic microenvironment present *in vivo*. By contrast, humanized mouse models provide an *in vivo* setting and have yielded valuable insights into HIV-1 pathogenesis, responses to ART and cure strategies^13,14^. However, commonly used bone marrow–liver–thymus (BLT) mice and related models develop poorly organized SLOs and depend on fetal tissue^15^, which is subjected to ethical and regulatory constraints.

A more physiologically relevant human system is therefore needed for HIV-1 research. Recent studies demonstrate that the lymphoid organ-chip, a microfluidic system in which high-density human PBMCs are embedded in a 3D extracellular matrix (ECM), can functionally recapitulate key features of human lymphoid tissue. In the two-channel chip originally developed by Goyal et al.^16^, primary human T and B cells under continuous perfusion autonomously assemble into germinal center-like lymphoid follicles (LFs) that contain activation-induced cytidine deaminase (AID)-expressing lymphocytes and that generate plasma cells and strain-specific antibodies following influenza vaccination. A subsequent study using a lymphoid organ-chip seeded with human PBMCs reproduced memory B cell responses to SARS-CoV-2 mRNA vaccine boosting^17^. Most recently, coupling the lymphoid organ-chip to a muscle module recapitulated immune responses to intramuscular mRNA vaccination, including somatic hypermutation and induction of neutralizing antibodies during recall responses^18^. Importantly, relative to static 3D culture, perfusion within the microfluidic chip improves lymphoid follicle formation, prevents B cells from autoactivation and non-specific IgG production, preserves the original B cell subset composition, and is required for spike-specific B cell responses elicited by SARS-CoV-2 mRNA vaccine boosting^16,17^.

The successful recapitulation of vaccine responses by the microfluidic lymphoid organ-chip prompted us to ask whether the microenvironment provided by this system can support natural HIV-1 replication and recapitulate key features of HIV-1 immunopathogenesis. While microfluidic devices have been applied to HIV-1 diagnosis and monitoring^19^, to our knowledge, the organ-chip technology, in which perfused microfluidic culture is used to reproduce tissue-level structure and function, has not been used to model HIV-1 infection^20^. Here we show that both X4 and R5 tropic HIV-1 strains replicate productively in the lymphoid organ-chip without exogenous T cell stimulation or cytokine supplementation. Infection recapitulates several hallmarks of HIV-1 immunopathogenesis, including CD4 T cell depletion, a decreased CD4/CD8 ratio, loss of T follicular helper (Tfh) cells, and altered T cell turnover and activation. ART applied to infected chips suppresses HIV-1 replication, reduces HIV-1 reservoir size, protects CD4 T cells, and reverses T cell phenotype alterations induced by HIV-1 infection. The productive infection of the lymphoid organ-chip by diverse HIV-1 strains establishes proof of concept that this platform can serve as a physiologically relevant, immunocompetent human model for both fundamental and translational HIV-1 research.

## Results

### Lymphoid organ-chip largely preserves *in vivo* T cell subset diversity

Previous studies have demonstrated that the lymphoid organ-chip can functionally recapitulate key features of human secondary lymphoid organs (SLOs) in terms of B cell responses to vaccination^16–18^. Because SLOs are the principal sites at which HIV-1 replicates and persists *in vivo*, we sought to determine whether the lymphoid organ-chip can support HIV-1 replication without exogenous T cell activation and recapitulate key features of HIV-1 immunopathogenesis (Fig. 1a).

**Fig. 1.**
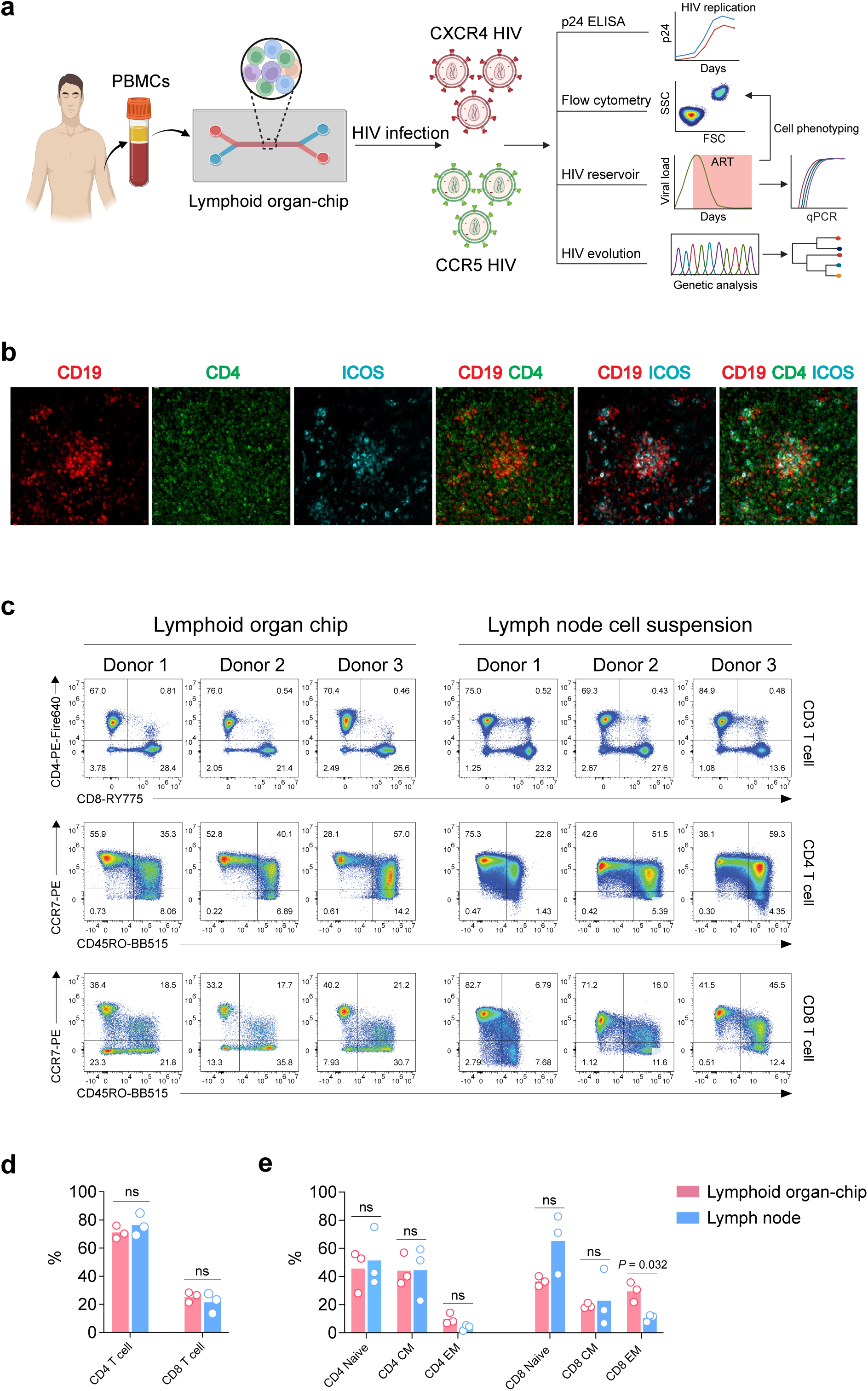
The lymphoid organ-chip forms ectopic lymphoid follicles and preserves the T cell subset composition of human lymph node. **a**, Schematic of the overall study design. Human PBMCs were seeded into a two-channel microfluidic lymphoid organ-chip and infected with either X4 or R5 tropic HIV-1. Viral replication kinetics, cell phenotyping, integrated proviral DNA were determined in the absence or presence of ART. Viral evolution in the chip was determined by single genome amplification (SGA). **b**, Representative immunofluorescence images of a lymphoid follicle formed in the chip at day 4 after cell seeding, stained for CD19 (red), CD4 (green) and ICOS (cyan). **c**, Flow cytometry plots showing T cell phenotypes recovered from lymphoid organ-chips (left) and from human lymph node cell suspensions (right). Three donors were studied per group. For lymphoid organ-chip, cell phenotype was determined five days after cell seeding. Top, CD4 and CD8 expression gated on CD3⁺ T cells; middle and bottom, CCR7 and CD45RO expression gated on CD4⁺ and CD8⁺ T cells, respectively, defining naïve (CCR7⁺CD45RO⁻), central memory (CM; CCR7⁺CD45RO⁺) and effector memory (EM; CCR7⁻CD45RO⁺) subsets. Numbers indicate the percentage of cells in each quadrant. **d**, Percentages of CD4 and CD8 T cells among CD3⁺ T cells in lymphoid organ-chips (pink) and lymph node (blue). **e**, Percentages of naïve, CM and EM subsets within the CD4 and CD8 T cell compartments. Bars represent mean of each group. Symbols represent individual donors (n = 3 per group). Statistical difference was determined by unpaired *t*-test; ns, not significant.

As previously reported, lymphoid follicle (LF) formation was observed four days after cell seeding (Fig. 1b). B cells formed clear clusters together with ICOS⁺ CD4 T cells (Fig. 1b), confirming that B and T cells autonomously assemble into ectopic LFs on the chip without exogenous stimulation. Previous studies also showed that the chip maintained natural B cell subset diversity without autoactivation, whereas autoactivation and non-specific IgG production occurred under static 3D culture conditions^16^. Because the cells most relevant to HIV-1 infection are T cells, and CD4 T cells in particular, we next determined whether the lymphoid organ-chip also maintains the T cell subset diversity observed *in vivo*. We recovered cells from the chip five days after seeding and compared their phenotype with that of cells recovered from lymph node samples from three donors. T cells recovered from the chips showed frequencies of CD4 and CD8 T cells comparable to those in lymph node (Fig. 1c–d). We further compared the frequency of each T cell subset — naïve, central memory (CM) and effector memory (EM) — between chip and lymph node. In the CD4 compartment, there was no significant difference in the frequency of any subset between the two sources (Fig. 1c–d). In the CD8 compartment, the only significant difference was in the EM subset, with cells recovered from the chip showing a higher EM frequency than lymph node cells (Fig. 1c–d), suggesting that CD8 T cells in the chip may be more activated than *in vivo*. Overall, however, the lymphoid organ-chip largely preserves the *in vivo* T cell subset composition, particularly for CD4 T cells.

### Both CXCR4 and CCR5 tropic HIV-1 strains replicate productively in the lymphoid organ-chip without artificial T cell activation

Productive HIV-1 replication in 2D culture requires exogenous T cell activation, most commonly with PHA or anti-CD3/CD28 antibodies. A major disadvantage of artificial stimulation is that T cells lose the subset diversity they display *in vivo*: after three days of PHA or anti-CD3/CD28 stimulation, which is routinely performed before HIV-1 infection, the vast majority of CD4 T cells exhibited a CCR7⁺ central memory phenotype (Extended Data Fig. 1a). By contrast, CD4 T cells in HIV infected individuals maintain clear subset diversity during both acute and chronic infection (Extended Data Fig. 1b). Because CD4 T cell subset diversity is well preserved in the lymphoid organ-chip, we sought to determine whether HIV-1 can replicate productively in the chip without exogenous stimulation. To this end, chips were infected with six HIV-1 strains, including three X4 tropic (40700, NL4.3, SG3) and three R5 tropic viruses (CH058, CH040, TRJO). To determine HIV-1 replication kinetics, chips were infected with each virus (5 ng p24 per 10^6^ cells, approximately 0.001 MOI) at day 4 after cell seeding. We chose this time point because it is the earliest at which an *in vivo*-like lymphoid structure has formed in the chip (Fig. 1b). We used the same amount of viral input as in our previous 2D culture studies^21^. To compare viral replication kinetics between the chip and static 3D culture, cells resuspended in the same ECM gel were infected with the same dose of each virus and cultured under static conditions. HIV-1 replication kinetics were monitored by measuring p24 concentration in the chip outflow or, for static cultures, in the supernatant. All six viruses replicated efficiently in the chip in all three PBMC donors tested, as reflected by exponentially increasing p24 concentrations (Fig. 2a). Peak p24 production occurred between day 8 and day 14 after infection, depending on both virus strain and PBMC donor (Fig. 2a). Under static 3D conditions, by contrast, only three viruses (NL4.3, CH040 and TRJO) replicated consistently across all three donors, and with delayed kinetics relative to the microfluidic condition (Fig. 2a). Notably, the X4 tropic transmitted/founder (T/F) virus 40700 failed to replicate under static conditions in any of the three donors, whereas SG3 and CH058 replicated in two of three donors (Fig. 2a). These data indicate that the lymphoid organ-chip supports productive replication of both R5 and X4 tropic HIV-1 without the need for exogenous T cell activation or cytokine supplementation, and that static 3D culture in the same ECM gel supported productive replication less consistently.

**Fig. 2.**
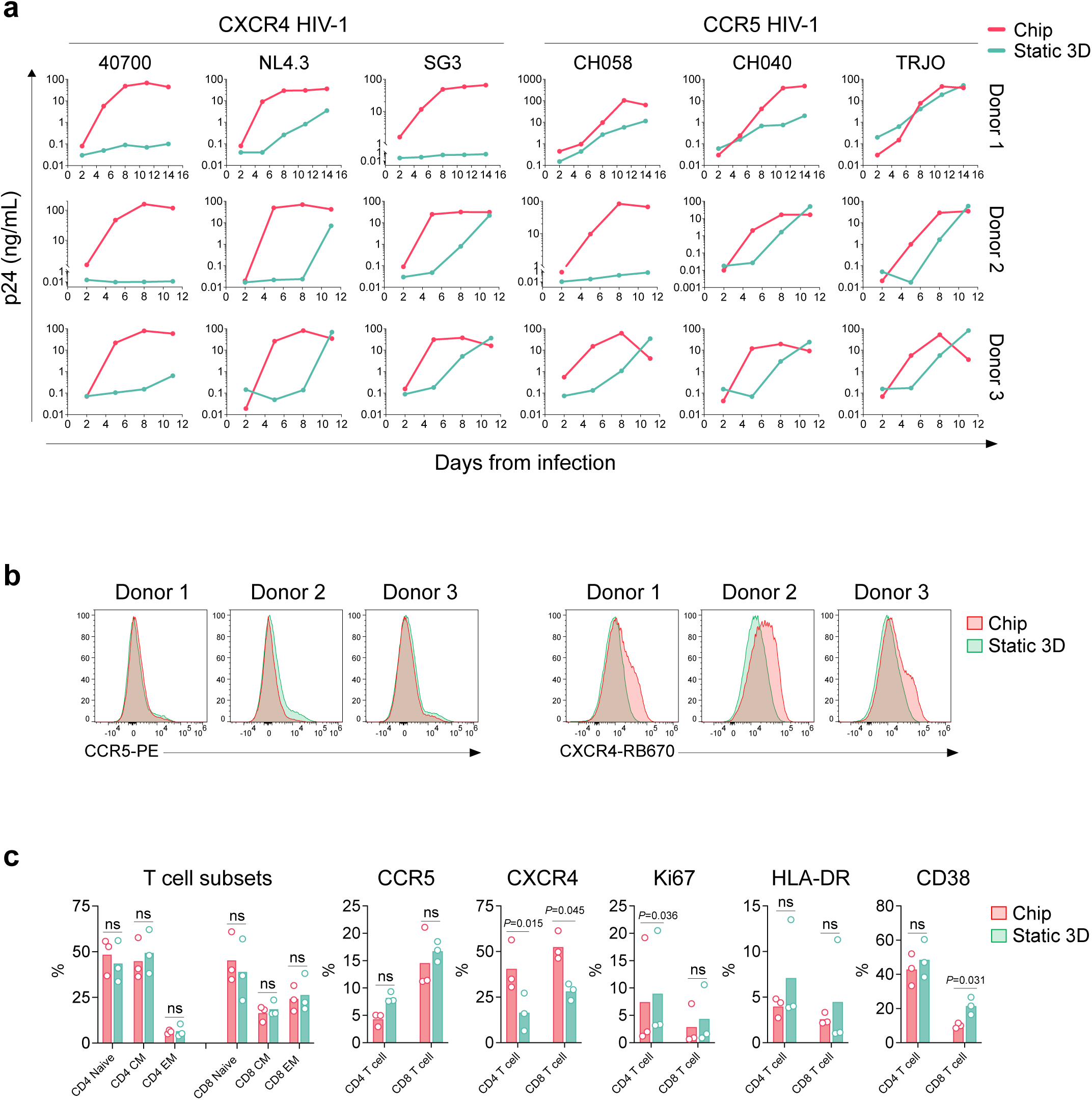
Both X4 and R5 tropic HIV-1 strains replicate productively in lymphoid organ-chip without exogenous T cell activation, which is more permissive than matched static 3D culture. **a**, HIV-1 replication kinetics in lymphoid organ-chips (pink) and matched static 3D cultures (teal) after infection with three X4-tropic (40700, NL4.3, SG3) and three R5-tropic (CH058, CH040, TRJO) HIV-1 strains. Each row shows one PBMC donor. p24 concentration was measured in chip outflow or static culture supernatant at the indicated times after infection and is plotted on a logarithmic scale. **b**, Flow cytometry histograms showing CCR5 (left) and CXCR4 (right) expression on T cells recovered from uninfected lymphoid organ-chips (pink) and uninfected static 3D cultures (teal) at day 4 after cell seeding in three donors. Histograms are normalized to mode. **c**, Frequencies of naïve, central memory (CM) and effector memory (EM) CD4 and CD8 T cell subsets, and expression of CCR5, CXCR4, Ki67, HLA-DR and CD38 on CD4 and CD8 T cells, in lymphoid organ-chips (pink) and static 3D cultures (teal). Bars show mean of each group. Symbols represent in individual donors (n = 3). Statistical difference was determined using paired *t*-test; ns: not significant.

To confirm that HIV-1 replication in the chip is indeed productive, we further determined viral replication kinetics by quantifying HIV-1 RNA viral load (VL) and infectivity using one X4 virus (40700) and one R5 virus (TRJO). As with the kinetics determined by p24 concentration, both HIV-1 RNA VL and infectivity increased exponentially (Extended Data Fig. 2). Viral infectivity, determined as relative luciferase units (RLU) in TZM-bl cells, increased in parallel with HIV-1 RNA VL, and both peaked on day 6 for 40700 and day 8 for TRJO (Extended Data Fig. 2). These results confirm that virions released from the chip are infectious and that the exponential increase in p24 and HIV-1 RNA represents productive HIV-1 replication. We also asked whether a cytokine cocktail containing IL-7, IL-15, TNF-α, IL-6 and IL-10 (25 ng/mL each), which promotes CD4 T cell proliferation and differentiation^22^, could enhance HIV-1 replication in the chip. Replication in the presence of the cocktail yielded higher p24 concentrations for all six strains (Extended Data Fig. 3a). Finally, to determine whether cells recovered from human lymph node support replication better than PBMCs, we seeded chips with lymph node cell suspensions from one donor and infected them with one X4 (40700) and one R5 (CH058) strain. HIV-1 replicated less efficiently in these chips than in those seeded with PBMCs (Extended Data Fig. 3b).

The difference in replication kinetics between the chip and static 3D culture suggested that CD4 T cell phenotype might differ between the two conditions. We therefore compared the baseline (uninfected) T cell phenotype between the two conditions. Static 3D culture showed slightly increased CCR5 expression and markedly decreased CXCR4 expression relative to the chip (Fig. 2b). Because CCR5 expression increases during T cell activation whereas CXCR4 expression decreases, this suggested that T cells are more activated under static 3D conditions, as previously observed for B cells^16^. To test this, we determined the frequency of each T cell subset together with the expression of proliferation and activation markers. There was no significant difference in T cell subset frequencies between the two conditions (Fig. 2c). However, T cells in static 3D culture tended to show increased expression of the proliferation marker Ki67 and the activation markers HLA-DR and CD38, in addition to CCR5 (Fig. 2c). The increases in Ki67 on CD4 T cells and CD38 on CD8 T cells were statistically significant, and the decrease in CXCR4 expression under static 3D conditions was significant in both CD4 and CD8 T cells (Fig. 2c). Together, these data indicate that, as previously observed for B cells, T cells in static 3D culture tend to be more activated than in the microfluidic chip. However, because T cell activation is generally considered to favor HIV-1 replication, these data suggest that factors other than the level of T cell activation are likely to underlie the higher permissiveness of the microfluidic chip.

### The lymphoid organ-chip recapitulates HIV-1 induced CD4 T cell depletion and the decreased CD4/CD8 ratio

The successful replication of both X4 and R5 HIV-1 strains in the chip allowed us to ask whether key features of HIV-1 immunopathogenesis could be recapitulated in this system. We first determined CD4 and CD8 T cell dynamics at the subset level. Relative to uninfected control chips, all six HIV-1 strains caused CD4 T cell depletion in all three PBMC donors (Fig. 3a). All six significantly depleted EM CD4 T cells, and five of six (except CH040) significantly depleted CM CD4 T cells. Naïve CD4 T cell depletion was highly variable and appeared more pronounced for the X4 strains (Fig. 3a). For CD8 T cells, three of six viruses — two X4 strains (40700 and NL4.3) and one R5 strain (TRJO) — caused consistent and significant CD8 T cell expansion in all three donors. Expansion was also observed with the remaining three viruses in some donors, particularly in the EM CD8 subset (Fig. 3a).

**Fig. 3.**
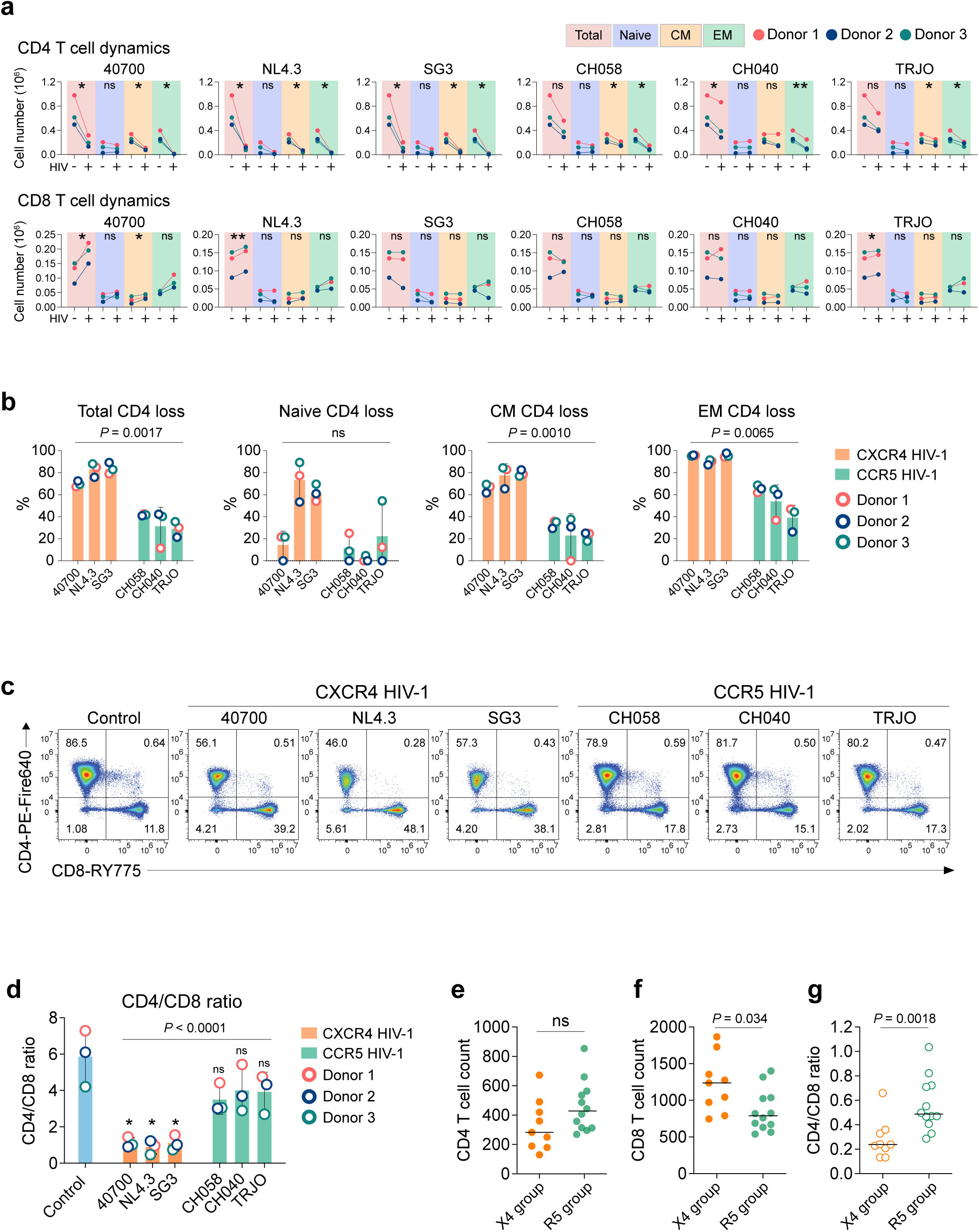
The lymphoid organ-chip recapitulates CD4 T cell depletion, the decreased CD4/CD8 ratio and the greater pathogenicity of X4 tropic HIV-1. **a**, Absolute numbers of total, naïve, central memory (CM) and effector memory (EM) CD4 (top) and CD8 (bottom) T cells recovered from uninfected (−) and HIV-1 infected (+) lymphoid organ-chips at day 9 after infection. Different T cell subsets are color-coded with background shading. Symbols denote individual PBMC donors, with lines connecting paired uninfected and infected chips from the same donor. Statistical difference was determined using paired *t*-test. \**P* < 0.05, \*\**P* < 0.01; ns, not significant. **b**, Loss of total, naïve, CM and EM CD4 T cells, expressed as the percentage reduction relative to uninfected control chips, for three X4 and three R5 HIV-1 strains. Statistical difference between X4 and R5 groups was determined using unpaired *t*-test, ns, not significant. **c**, Representative flow cytometry plots of CD4 and CD8 expression gated on CD3⁺ T cells from an uninfected control chip and from chips infected with each of the six HIV-1 strains (donor 1, day 9 after infection). **d**, CD4/CD8 ratio in uninfected control chips and in chips infected with each HIV-1 strain from three different donors. Statistical difference between X4 and R5 groups was determined using unpaired *t*-test. Statistical difference between infected and uninfected chips was determined using paired *t*-test. \**P* < 0.05; ns, not significant. **e-g**, CD4 T cell count (**e**), CD8 T cell count (**f**) and CD4/CD8 ratio (**g**) in 21 participants from the RV217 Thailand cohort harboring X4 variants (n = 9) or only R5 variants (n = 12) at around two years after HIV-1 transmission. Horizontal line represents the mean. Statistical difference between groups was determined using two-tailed Mann–Whitney U test; ns: not significant.

X4 tropic HIV-1 is more pathogenic than R5 tropic HIV-1 and causes faster CD4 T cell depletion and disease progression^6,23,24^. We previously showed that X4 viruses preferentially target and deplete the CM CD4 subset *in vivo*^2,25^. We therefore compared the degree of CD4 T cell depletion between X4 and R5 strains to determine whether the chip recapitulates this difference in pathogenesis. X4 strains caused significantly greater total CD4 T cell depletion than R5 strains (Fig. 3b). The difference was most apparent in the CM CD4 subset and was also statistically significant in the EM CD4 subset (Fig. 3b).

Another hallmark of HIV-1 pathogenesis is a decreased CD4/CD8 ratio. Relative to uninfected control chips, all six HIV-1 strains reduced the CD4/CD8 ratio in all three PBMC donors (Fig. 3c,d); the decrease reached statistical significance for all three X4 strains but not for the R5 strains (Fig. 3d). X4 infection also resulted in a significantly lower CD4/CD8 ratio than R5 infection (Fig. 3d). To determine whether this difference is also apparent in natural infection, we analyzed CD4 and CD8 T cell counts and the CD4/CD8 ratio in 21 participants of the RV217 Thailand cohort^26^, in whom we had previously determined coreceptor usage approximately two years after HIV-1 transmission and identified 9 harbored X4 variants and 12 harbored only R5 variants^25^. Participants harboring X4 variants had lower CD4 counts, significantly higher CD8 counts and a significantly lower CD4/CD8 ratio than those harboring only R5 variants (Fig. 3e–g). Together, these data show that the lymphoid organ-chip recapitulates not only the decreased CD4/CD8 ratio of HIV-1 infection but also the differential impact of X4 and R5 strains on this ratio.

In contrast, CD4 T cell depletion was not consistently observed under static 3D conditions. Of the six viruses, only SG3 consistently caused CD4 depletion in both donors analyzed (Extended Data Fig. 4a). Strain 40700, which failed to replicate under static conditions in any donor (Fig. 2a), was associated with increased CD4 counts relative to uninfected controls (Extended Data Fig. 4a). NL4.3, CH058, CH040 and TRJO replicated in both donors (Fig. 2a), yet CD4 T cell depletion remained inconsistent: TRJO was associated with increased CD4 counts in both donors, whereas NL4.3, CH058 and CH040 each caused depletion in only one donor (Extended Data Fig. 4a). All six viruses caused CD8 T cell expansion in both donors (Extended Data Fig. 4a). The decreased CD4/CD8 ratio under static conditions was therefore driven mainly by CD8 expansion, and no difference in CD4/CD8 ratio was observed between X4 and R5 strains (Extended Data Fig. 4b). Thus, unlike the lymphoid organ-chip, static 3D culture did not faithfully recapitulate CD4 depletion and the differential pathogenesis of X4 and R5 strains.

### HIV-1 infection induces loss of T follicular helper CD4 T cells in the lymphoid organ-chip

Extensive infection of Tfh cells in lymphoid tissue has been demonstrated during HIV-1 infection and during pathogenic SIV infection of rhesus macaques, whereas non-pathogenic SIV infection of natural hosts such as sooty mangabeys is associated with a significantly lower level of Tfh infection^27,28^. We therefore asked whether HIV-1 infection causes Tfh cell depletion in the lymphoid organ-chip. Analysis of CXCR5⁺PD-1⁺ Tfh cell dynamics revealed significant Tfh depletion by all six HIV-1 strains relative to uninfected control chips (Fig. 4a,b). Depletion was more pronounced for the X4 strains, which eliminated Tfh cells almost completely by day 9 after infection (Fig. 4a–c). For both X4 and R5 strains, the decline in CXCR5⁺PD-1⁺ cells was accompanied by a reduction in PD-1⁺ but not CXCR5⁺ CD4 T cells (Fig. 4a).

**Fig. 4.**
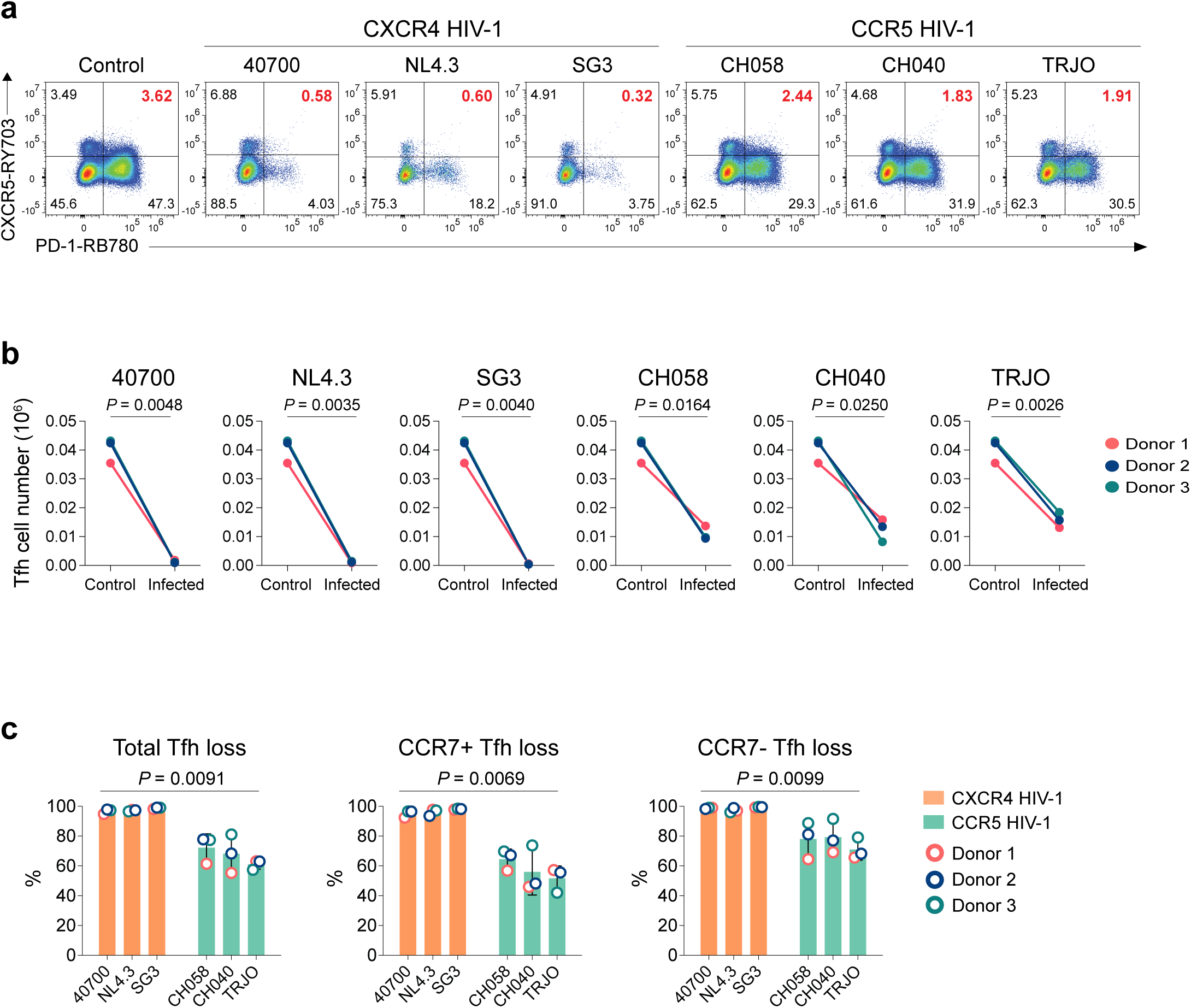
HIV-1 infection depletes T follicular helper cells in the lymphoid organ-chip. **a**, Representative flow cytometry plots of CXCR5 and PD-1 expression gated on CD4 T cells recovered from an uninfected control chip and from chips infected with each of six HIV-1 strains at day 9 after infection (donor 1). Red values denote the CXCR5⁺PD-1⁺ Tfh population. **b**, Absolute Tfh cell numbers in paired uninfected control and infected chips for each HIV-1 strain. Symbols denote individual PBMC donors, with lines connecting paired chips from the same donor. Statistical difference was determined using paired *t*-test. **c**, Loss of total Tfh, CCR7⁺ and CCR7⁻ Tfh cells, expressed as the percentage reduction compared to uninfected control chips, for three X4 and three R5 HIV-1 strains. Bars represent mean ± SD. Symbols show individual donors (n = 3). Statistical difference between X4 and R5 groups was determined using unpaired *t*-test.

Tfh cells in the chip were present in both the CCR7⁺ (CM) and CCR7⁻ (EM) CD4 compartments. Previous studies show that Tfh cells lose CCR7 on entering secondary lymphoid tissue, which permits their localization to B cell follicles and germinal centers, whereas CCR7 expression retains activated T cells in the T cell zone^29,30^. Analysis of each compartment showed that X4 strains depleted Tfh cells almost completely in both the CCR7⁺ and CCR7⁻ compartments, whereas R5 strains tended to deplete more Tfh cells in the CCR7⁻ compartment (Fig. 4c). This difference may reflect the higher CCR5 expression on CCR7⁻ (effector memory) CD4 T cells that we have previously described^25^.

CXCR5-expressing follicular cells with cytotoxic potential are also present in the CD8 compartment^31–33^. These follicular cytotoxic CD8 T (Tfc) cells may exert anti-HIV activity by killing infected Tfh cells within the germinal center^33^. We therefore also determined the impact of HIV-1 infection on Tfc CD8 T cells. We observed a trend towards Tfc loss with five of the six viruses in all three donors, while CH040 was associated with a slightly increased Tfc cell number in one donor (Extended Data Fig. 5a,b). However, unlike Tfh CD4 T cells, the extent of Tfc loss was highly variable across donors and did not reach statistical significance (Extended Data Fig. 5b), and there was no significant difference between X4 and R5 strains (Extended Data Fig. 5c). Because CD8 T cells could not be directly targeted by HIV-1, the loss of Tfc CD8 T cells is likely mediated by indirect mechanisms, such as immune dysfunction induced by HIV-1 infection.

### Divergent impacts of CXCR4 and CCR5 tropic HIV-1 on T cell turnover and activation in the lymphoid organ-chip

An important hallmark of HIV-1 pathogenesis is increased T cell turnover and immune activation, and the level of immune activation predicts disease progression better than set-point viral load^34–36^. Nonetheless, the principal cause of immune activation in natural HIV-1 infection remains controversial. While some studies support HIV-1 antigens as the major driver^37^, others implicate microbial translocation^38^. Because secondary lymphoid organs are the major sites of immune activation *in vivo*, the successful replication of HIV-1 in the lymphoid organ-chip without artificial T cell activation provides a unique opportunity to determine whether HIV-1 replication alone can elevate T cell turnover and activation. We therefore determined expression of the turnover marker Ki67 and the activation markers HLA-DR and CD38 on CD4 and CD8 T cells at day 9 after infection, around the peak of viral replication. As in HIV-1-infected individuals, there was considerable variation in T cell turnover and activation across donors (Fig. 5a–d). Nevertheless, several clear patterns emerged. In the CD4 compartment, R5 strains tended to increase turnover and activation relative to uninfected control chips, more so in the EM than the CM subset (Fig. 5a,b). Of the three R5 viruses, TRJO increased CD4 turnover and activation in all three donors, CH058 in two of three, and CH040 increased turnover in one donor and activation in two. When the EM subset was analyzed alone, CH058 and TRJO consistently increased turnover in all three donors and CH040 in two of three. All three R5 viruses increased EM CD4 activation in all three donors, and the increases for CH040 and TRJO were significant (Fig. 5a,b). In contrast, X4 strains significantly decreased CD4 T cell turnover and activation, most clearly in the CM subset (Fig. 5a,b). Overall, R5 viruses induced significantly higher CD4 turnover and activation than X4 viruses, most apparent in the CM subset (Fig. 5a,b).

**Fig. 5.**
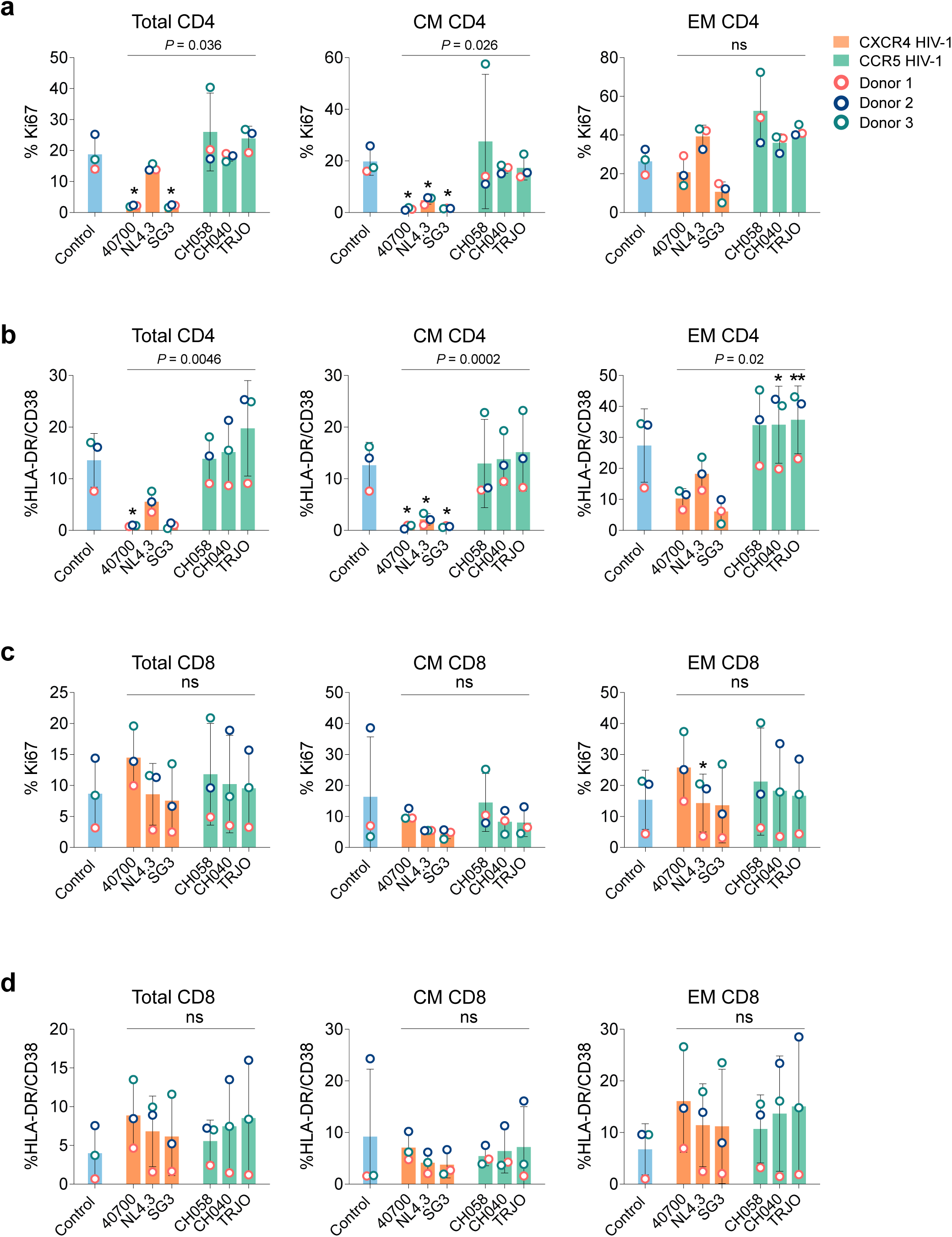
Impact of HIV-1 infection on CD4 and CD8 T cell turnover and activation in lymphoid-organ chip. **a,b**, Expression of Ki67 (**a**) and co-expression of HLA-DR and CD38 (**b**) in total, central memory (CM) and effector memory (EM) CD4 T cells recovered from uninfected control chips (blue) and from chips infected with three X4 (orange) or three R5 (green) HIV-1 strains at day 9 after infection. **c,d**, Corresponding analysis of Ki67 (**c**) and HLA-DR/CD38 (**d**) expression in total, CM and EM CD8 T cells. Bars show mean ± SD. Symbols represent individual PBMC donors (n = 3). Asterisks denote statistical comparison with the uninfected control chips using paired *t*-test; \**P* < 0.05, \*\**P* < 0.01. Statistical difference between R5 and X4 groups was determined using unpaired *t*-test; ns, not significant.

Because X4 HIV-1 caused significant CD4 depletion (Fig. 3a), we investigated whether this inhibition of cell turnover and activation reflected selective depletion of activated CD4 T cells. We therefore determined cell phenotype five days after X4 virus infection, before extensive CD4 depletion occurred. At this earlier time point, 30–44% of CD4 T cells were depleted relative to uninfected controls (Extended Data Fig. 6a). However, for each virus the decrease in CD4 T cell turnover and activation was approximately twofold greater than the extent of CD4 T cell depletion (Extended Data Fig. 6a). By contrast, turnover and activation of CD8 T cells tended to increase following X4 infection (Extended Data Fig. 6b). Together, these observations indicate that the inhibition of CD4 T cell turnover and activation by X4 viruses cannot be fully explained by direct CD4 T cell depletion, and that additional, yet unidentified mechanisms are likely to contribute.

The effect of HIV-1 infection on CD8 turnover was highly variable across viruses and donors (Fig. 5c). The effect on CD8 activation was more consistent: all viruses except SG3 and CH058 increased CD8 activation in all three donors, and SG3 and CH058 each decreased CD8 activation in a single donor (Fig. 5d). The increase in CD8 activation was more pronounced in EM than in CM CD8 T cells (Fig. 5d). There was no significant difference between X4 and R5 strains in inducing CD8 T cell turnover or activation (Fig. 5c,d).

To determine whether the level of immune activation induced in the chip reflects that of natural infection, we compared the frequencies of HLA-DR⁺CD38⁺ CD4 and CD8 T cells with frequencies previously reported in PBMC samples from acutely HIV-1-infected individuals in an acute infection cohort^39^. We restricted this analysis to R5 strains, since the vast majority of acute HIV-1 infections are established by R5 virus. Across all three donors, the median frequency of HLA-DR⁺CD38⁺ CD4 T cells induced by R5 infection in the chip was 15.5%, compared with reported frequencies of 2.1% in Fiebig stage I and 2.6% in Fiebig stages II–IV^39^. The median frequency of HLA-DR⁺CD38⁺ CD8 T cells induced by R5 infection in the chip was 7.2%, compared with 7.6% in Fiebig stage I and 14.9% in Fiebig stages II–IV^39^. Therefore, CD4 T cell activation in the chip exceeds that reported in acute HIV-1 infection, whereas CD8 activation is comparable to Fiebig stage I but lower than Fiebig stages II–IV. However, these data may not be directly comparable, since the chip models the lymph node microenvironment whereas the published frequencies derive from PBMC^39^, and T cell activation is higher in lymph node than in matched peripheral blood in HIV-1-infected individuals^40^. Altogether, the finding that HIV-1 replication in the chip increases T cell turnover and activation indicates that HIV-1 replication alone is sufficient to drive immune activation, supporting the view that immune activation in HIV-1 infection is driven, at least in part, by HIV-1 itself rather than requiring microbial translocation.

### ART suppresses HIV-1 replication and reduces integrated proviral DNA in the lymphoid organ-chip

We next determined whether antiretroviral therapy (ART) can suppress HIV-1 replication and reduce the size of HIV-1 reservoir, as occurs in HIV-1-infected individuals on ART (Fig. 6a)^41^. Chips were infected with one R5 (TRJO) and one X4 (40700) strain. These two viruses were chosen because their replication was completely inhibited by ART under static 3D conditions in preliminary experiments, excluding the possibility of drug resistance. Five days after infection, as viral replication approaching peak, chips were treated with a combination of three antiretroviral drugs: the protease inhibitor darunavir (10 μM), the integrase inhibitor raltegravir (10 μM) and the reverse transcriptase inhibitor nevirapine (10 μM). Infected chips without ART served as controls (Fig. 6a). In both R5 and X4 HIV-1 infected chips, HIV-1 RNA VL began to decrease two days after treatment (Fig. 6b), and by five days there were 199-fold and 175-fold decreases in VL in X4 and R5 HIV-1 infected chips, respectively, relative to untreated controls (Fig. 6b).

**Fig. 6.**
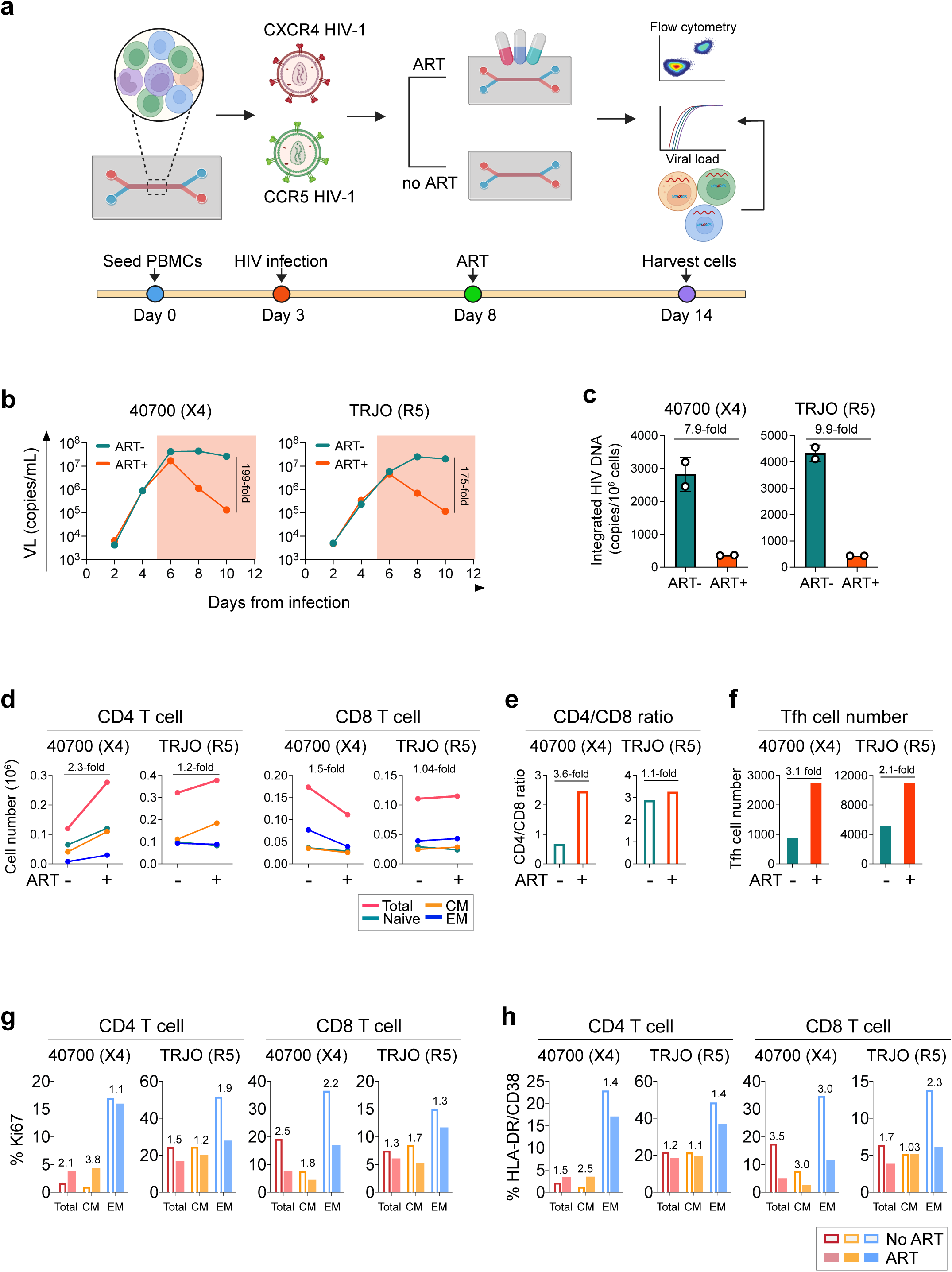
ART suppresses HIV-1 replication, reduces integrated proviral DNA and reverses infection-induced T cell phenotypic changes in the lymphoid organ-chip. **a**, Schematic and timeline of the experiment. Chips were infected with either X4 (40700) or R5 (TRJO) HIV-1 on day 3 after cell seeding. ART (darunavir, raltegravir and nevirapine; 10 μM each) was added 5 days after infection and cells were harvested on day 14 (6 days after ART initiation) for flow cytometry and quantification of integrated HIV-1 DNA. Viral load was monitored in chip outflow. **b**, Dynamics of HIV-1 RNA VL in chip outflow following infection by 40700 (left) or TRJO (right). Shading indicates the period of ART. The average VL of two independent VL assays is shown. Values denote fold reduction in VL relative to untreated chips five days after ART initiation. **c**, Integrated HIV-1 DNA per 10⁶ cells recovered from untreated and ART-treated chips. Values denote fold reduction. Mean and SD of two independent assays are shown. **d**, Absolute numbers of total, naïve, CM and EM CD4 (left) and CD8 (right) T cells in untreated and ART-treated chips. Values show fold change in total T cell number. **e,f**, CD4/CD8 ratio (**e**) and absolute Tfh cell number (**f**) in untreated and ART treated chips. Values denote fold changes. **g,h**, Ki67 expression (**g**) and HLA-DR/CD38 co-expression (**h**) in total, CM and EM CD4 (left) and CD8 (right) T cells from chips without ART (open bars) or with ART (filled bars). Values denote fold changes between treated and untreated conditions.

To assess the impact of ART on HIV-1 reservoir size, we recovered cells from infected chips six days after treatment and quantified integrated HIV-1 DNA. Compared with untreated chips, integrated HIV-1 DNA was reduced 7.9-fold in X4 HIV-1 infected chips and 9.9-fold in R5 HIV-1 infected chips (Fig. 6c). Together, these data demonstrate that applying ART to HIV-1 infected chips not only rapidly inhibits HIV-1 replication but also substantially reduces the burden of integrated provirus.

### ART preserves CD4 T cells and reverses the T cell activation phenotype induced by HIV-1 infection

Given the rapid inhibition of HIV-1 replication by ART in the chip, we further determined its impact on T cell phenotype. Six days after treatment, CD4 T cell counts increased 2.3-fold in X4 HIV-1 infected chips and 1.2-fold in R5 HIV-1 infected chips relative to untreated controls (Fig. 6d). In X4 virus infected chips, increased CD4 T cell numbers were observed in all three subsets, whereas in R5 virus infected chips the increase was confined to the CM subset (Fig. 6d). In X4 virus infected chips, six days of ART also led to a 1.5-fold decrease in CD8 T cell count, which was not observed in R5 virus chips (Fig. 6d). Accordingly, ART resulted in a marked 3.6-fold increase in CD4/CD8 ratio in X4 HIV-1 chip and a slight 1.1-fold increase in R5 HIV-1 infected chip (Fig. 6e), consistent with the lower CD4/CD8 ratio caused by X4 infection. Notably, ART also protected Tfh CD4 T cells: six days of treatment resulted in 3.1-fold and 2.1-fold increases in Tfh numbers in X4 and R5 HIV-1 infected chips, respectively (Fig. 6f). These data demonstrate that applying ART to HIV-1 infected lymphoid organ-chip protects CD4 T cells, including Tfh cells, from HIV-1 induced cell depletion.

We next determined the impact of ART on HIV-1-induced T cell proliferation and activation. In particular, we asked whether this impact differs between X4 and R5 HIV-1 infection, given their opposite effects on CD4 T cell proliferation and activation (Fig. 5a–b). In X4 HIV-1 infected chips, ART increased the proliferation and activation of total and CM CD4 T cells relative to untreated controls (Fig. 6g-h), consistent with the inhibition of CD4 proliferation and activation by X4 HIV-1, particularly in the CM CD4 subset (Fig. 5a-b). By contrast, in R5 HIV-1 infected chips, ART decreased CD4 proliferation and activation relative to untreated controls, most obviously in the EM CD4 subset (Fig. 6g-h), consistent with the increase in EM CD4 proliferation and activation caused by R5 HIV-1 (Fig. 5a-b). For CD8 T cells, ART decreased cell proliferation and activation in both X4 and R5 HIV-1 infected chips (Fig. 6g-h), consistent with the increase in CD8 activation caused by both X4 and R5 HIV-1 strains (Fig. 5c-d). In summary, these observations demonstrate that suppressing HIV-1 replication with ART reverses the T cell phenotypic alterations induced by HIV-1 infection and confirm that the effects of HIV-1 infection on CD4 and CD8 T cell proliferation and activation, including the inhibition of CD4 T cells proliferation and activation by X4 HIV-1, are driven mainly by viral replication.

### HIV-1 evolution in the lymphoid organ-chip

Productive HIV-1 replication in the lymphoid organ-chip implies that the virus should evolve over time. We therefore investigated the pattern of HIV-1 evolution in a CH058 infected chip two weeks after infection by sequencing the full-length envelope region using single genome amplification (SGA). Phylogenetic tree and Highlighter plot analysis of 11 envelope SGA sequences showed that the virus diversified from the original (ancestral) sequence through the accumulation of point mutations (Fig. 7a), and the consensus of the 11 sequences exactly matched the ancestral sequence (Fig. 7a). To further characterize the evolutionary pattern, we analyzed the sequences using the Poisson Fitter tool in the Los Alamos HIV sequence database. The results showed that the sequences followed a star-like phylogeny, and Hamming distances followed a Poisson distribution (Fig. 7b), indicating that HIV-1 evolved in the lymphoid organ-chip without detectable selection pressures during the first 14 days.

**Fig. 7.**
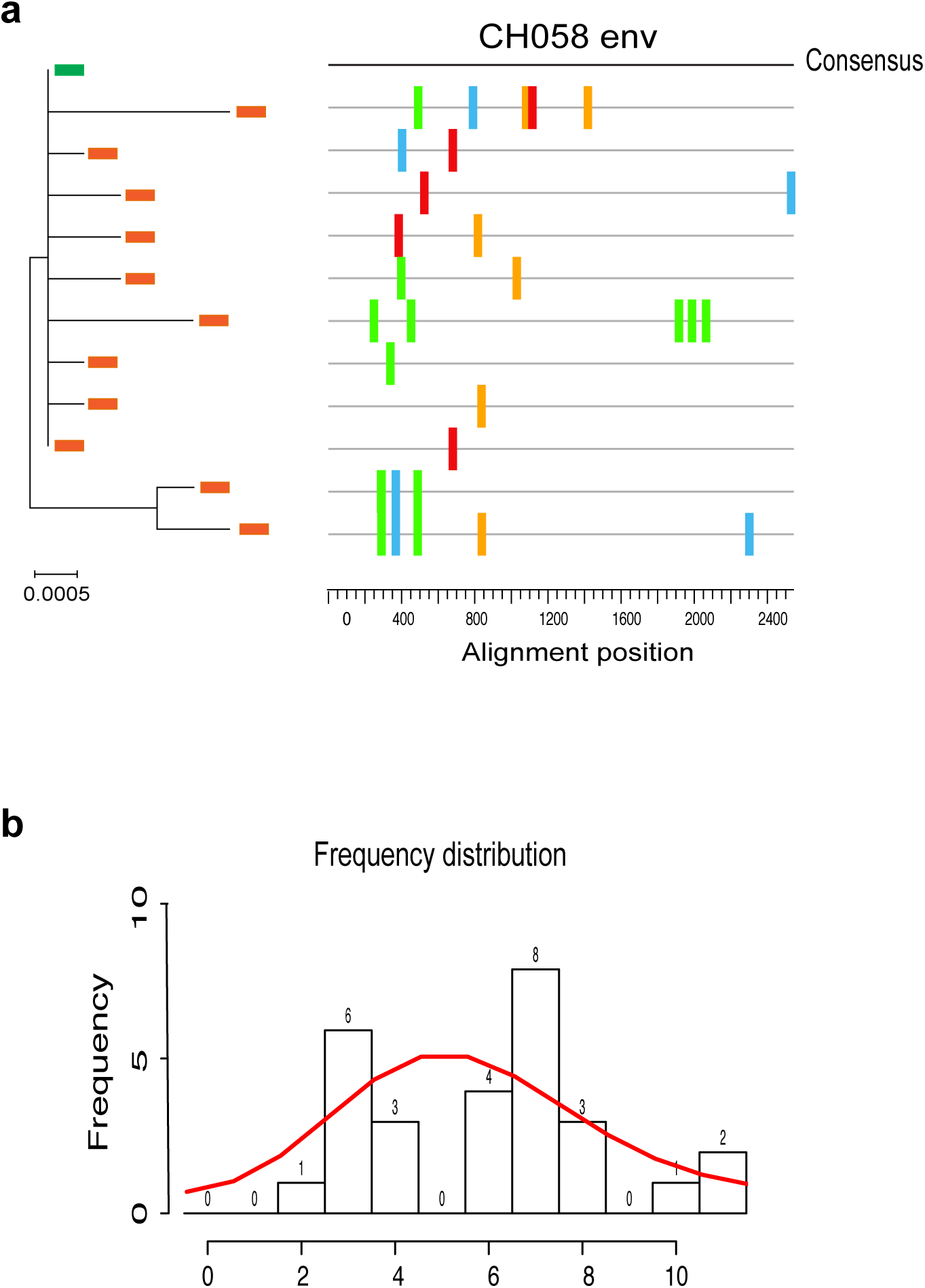
HIV-1 evolution in the lymphoid organ-chip. **a**, Maximum-likelihood phylogenetic tree (left) and Highlighter plot (right) of 11 full-length HIV-1 *env* sequences obtained by single-genome amplification (SGA) from a CH058 infected lymphoid organ-chip at day 14 after infection (donor 1). In the Highlighter plot, each row represents one sequence and tick marks indicate nucleotide substitutions relative to the consensus sequence (top). **b**, Distribution of pairwise Hamming distances among the 11 sequences as determined using Poisson Fitter tool in Los Alamos HIV Sequence Database. The phylogenetic tree was constructed using Mega 6.

## Discussion

Here we demonstrate that a microfluidic lymphoid organ-chip supports productive, multi-round HIV-1 replication in the absence of exogenous T cell activation or cytokine supplementation, and that infection in this system recapitulates core features of HIV-1 immunopathogenesis at CD4 subset resolution. All six X4 and R5 HIV-1 strains tested replicated to peak titers within 8–14 days across three donors, whereas matched static 3D cultures supported consistent replication for only three viruses, indicating that perfusion, rather than the ECM or three-dimensionality alone, plays a major role in permissiveness. The chip preserved the CD4 T cell subset composition of human lymph node tissue, and infection recapitulated CD4 T cell depletion, a decreased CD4/CD8 ratio, loss of Tfh cells, and the greater pathogenicity of X4 strains, including their preferential depletion of central memory CD4 T cells as we previously showed in HIV-1-infected individuals^2,25^.

One question that remains to be addressed mechanistically is why the perfused microfluidic condition is more permissive for HIV-1 replication than static 3D culture. Phenotypic analysis showed that T cells tend to be more activated in static 3D culture than in the perfused organ-chip. This aligns with previous findings on the same platform, in which B cells under static conditions showed autoactivation without antigen stimulation, whereas the perfused chip retained the original B cell subset composition and prevented autoactivation^16^. However, because CD4 T cell activation is generally considered to favor HIV-1 replication, the activation phenotype itself cannot explain the difference in permissiveness between the microfluidic and static conditions. Several other factors may be at play. First, the replication kinetics of all six viruses indicate that the advantage of the microfluidic condition is greater for X4 viruses (Fig. 2a). This agrees with our finding that static 3D culture showed significantly decreased CXCR4 expression and slightly increased CCR5 expression (Fig. 2b,c). Indeed, these coreceptor expression differences further suggest that the microenvironment of the lymphoid organ-chip resembles that of the lymph node, since CXCR4 is expressed predominantly on naïve T cells, which are the principal lymph-node-homing subset, whereas CCR5 expression is largely confined to activated T cells that migrate through peripheral tissues^42^. Second, the environment within static 3D culture may restrict infection by cell-free virions while promoting cell-associated transmission, as shown in a previous study using collagen as a tissue-like 3D scaffold^43^. Third, improved nutrient and waste exchange, better sustained cell viability, and perfusion-induced changes in cellular metabolism and cytokine gradients may also increase permissiveness for HIV-1. Such factors could explain why the lymphoid organ-chip supported faster replication of R5 virus despite the lower CCR5 expression in the chip. A more comprehensive comparison of cell phenotype between the microfluidic chip and static 3D culture will be needed to define the mechanisms underlying the higher permissiveness of the microfluidic chip.

The preservation of *in vivo* T cell subset composition while supporting highly productive HIV-1 infection is a key advantage of the lymphoid organ-chip for HIV-1 research. It removes the requirement for exogenous T cell activation inherent to conventional 2D culture. In comparison with static tissue explants and organoid models, it requires only peripheral blood rather than surgical tissue. CD4 T cell subset identity is a critical parameter for understanding HIV-1 pathogenesis, reservoir persistence, transmission and immune reconstitution on ART. For pathogenesis, a low HIV-1 burden in the central memory CD4 T cell subset is a shared hallmark of both non-pathogenic SIV infection in natural hosts and durable, non-progressive HIV-1 infection in humans, although the mechanisms underlying remain an open question^44–48^. For reservoir persistence, the CM CD4 T cell subset has been identified as a major HIV-1 reservoir in people on ART^1^. In relation to HIV-1 transmissibility, our recent studies demonstrate that R5 HIV-1, which has a marked transmission advantage over X4 virus, has a replication advantage in the effector memory CD4 subset than X4 HIV-1^25^. Because EM CD4 T cells are located primarily at mucosal sites where most HIV-1 transmissions occur, this finding therefore suggests that HIV-1 replicative fitness in the EM CD4 subset may be an important determinant of HIV-1 transmissibility. Furthermore, an intact central memory CD4 compartment also predicts better immune reconstitution on ART^49^. Collectively, a deeper mechanistic understanding of these questions will provide critical insight into HIV-1 functional cure and prevention approaches. In this regard, the lymphoid organ-chip opens the possibility of investigating HIV-1 pathogenesis, transmissibility and reservoir at CD4 T cell subset resolution in a human-relevant, immunocompetent system.

HIV-1 infection in the lymphoid organ-chip not only recapitulated several fundamental features of natural HIV-1 immunopathogenesis, including CD4 T cell depletion, a decreased CD4/CD8 ratio and Tfh cell depletion, but also reproduced the differential pathogenicity of R5 and X4 HIV-1 strains. The significantly greater total CD4 T cell depletion by X4 HIV-1 in the chip is consistent with the long-standing observation that emergence of X4 virus through coreceptor switching is associated with faster CD4 T cell depletion and accelerated progression to AIDS^23–25^. The significantly greater depletion of the central memory CD4 subset by X4 HIV-1 is likewise consistent with our previous findings in natural HIV-1 infection that X4-tropic HIV-1 preferentially infects and depletes CM CD4 T cells compared with R5 HIV-1 in vivo^2,25^. Furthermore, the significantly lower CD4/CD8 ratio induced by X4 than by R5 virus in the chip was concordant with observations in 21 participants from the RV217 Thailand cohort. By contrast, static 3D culture did not consistently recapitulate CD4 T cell depletion, most likely owing to relatively poor HIV-1 replication. The decreased CD4/CD8 ratio under static conditions was driven mainly by CD8 T cell expansion, and no significant difference in CD4/CD8 ratio was observed between X4 and R5 strains. The lymphoid organ-chip therefore recapitulates the immunopathogenesis of both X4 and R5 HIV-1 more faithfully than static 3D culture in terms of CD4 and CD8 T cell dynamics.

Although elevated T cell turnover and immune activation are hallmarks of pathogenic HIV-1 infection, whether they are driven principally by HIV-1 replication and viral antigens or by indirect mechanisms such as microbial translocation remains controversial^34,35,38^. The lymphoid organ-chip offers a unique opportunity to address this question, because T cells are not artificially activated, and the system contains neither microbial translocation nor bacterial products. Our data therefore indicate that HIV-1 replication alone is sufficient to alter T cell turnover and immune activation, in a tropism-dependent manner (see below). When compared with acutely HIV-1-infected individuals, CD4 T cell activation induced by R5 virus in the chip was higher than that observed in PBMC during acute HIV-1 infection, whereas CD8 T cell activation was similar to that in Fiebig stage I but lower than in Fiebig stages II–IV^39^. Because lymph node T cell turnover and activation exceeds that in paired blood *in vivo*^40,50^, and the lymphoid organ-chip models the lymph node microenvironment, the higher CD4 T cell activation in the chip may reflect this compartmental difference, a possibility that will need to be confirmed by direct comparison with lymph node samples from HIV-1-infected individuals.

A finding not previously reported is the divergent effect of X4 and R5 HIV-1 on CD4 T cell turnover and activation. Whereas R5 HIV-1 increased immune activation in both the CD4 and CD8 compartments, most evidently in the EM subset, X4 HIV-1 specifically decreased turnover and activation of CD4 T cells, particularly in the CM subset, while increasing CD8 T cell activation. We further showed that this effect could not be fully explained by selective depletion of activated CD4 T cells by X4 viruses, as the inhibition was evident early after infection, before significant CD4 depletion, and was approximately twofold greater than the extent of CD4 depletion. Importantly, ART reversed both the inhibitory effect of X4 virus on CD4 T cell turnover and activation and the activating effect of R5 virus, indicating that both stem from active HIV-1 replication. Notably, this phenomenon would be difficult to identify in natural HIV-1 infection, since we have previously shown that even in people who have undergone coreceptor switching and harbor X4-tropic variants *in vivo*, R5 viruses continue to co-exist as the dominant population in plasma^25^. Defining the mechanism underlying X4 HIV-1 mediated inhibition of CD4 T cell turnover and activation will therefore be important, as it may represent a previously uncharacterized contributor to X4 HIV-1-induced pathogenesis. Because the effect is restricted to CD4 T cells, it is more likely to arise from a CD4-dependent mechanism related to direct infection than from an indirect mechanism such as inflammatory cytokine signaling.

The current study has several limitations. First, PBMC-derived lymphoid follicles represent ectopic lymphoid structures that lack the follicular dendritic cell (FDC) and fibroblastic reticular cell (FRC) networks, high endothelial venules, afferent lymph and cell trafficking of a true lymph node, leaving considerable room for further development through incorporation of these cell types and structures. Second, although previous work on this platform demonstrated germinal-center-like follicle formation, plasma cell differentiation and antigen-specific antibody responses following vaccination, we did not assess B cell responses induced by HIV-1 infection in the current study. The successful replication of diverse HIV-1 strains shown here, together with these earlier demonstrations of vaccine responsiveness, nonetheless positions the lymphoid organ-chip as a human system that can support both a functional vaccine response and productive HIV-1 infection in the same device, which opens the possibility of evaluating both preventive and therapeutic HIV vaccines. Third, although we show as proof of concept that ART applied to the chip rapidly inhibits HIV-1 replication, limits the reservoir and reverses HIV-1-induced T cell phenotypic alterations, we did not characterize reservoir dynamics under long-term ART or viral rebound following treatment interruption. A recent study using tonsil and spleen immune organoids has demonstrated ART-mediated suppression and viral rebound after analytical treatment interruption (ATI) with a subtype A HIV-1 strain^11^. Future studies will be required to determine viral load and reservoir dynamics under relatively long-term ART, as well as rebound upon ATI, in the perfused lymphoid organ-chip.

In summary, the productive replication of diverse HIV-1 strains in a perfused, immunocompetent human system without artificial activation, alongside the previously demonstrated capacity of this platform to support functional vaccine responses, establishes the lymphoid organ-chip as a model in which HIV-1 infection and host immunity can be interrogated together in the same human system, for both fundamental and translational HIV-1 research.

## Acknowledgements

We thank Iset Vera and the USF Health Microscopy and Cell Imaging Core for technical assistance. The authors acknowledge the Tampa General Hospital–University of South Florida Health Precision Medicine Biorepository (RRID:SCR_027501) for the collection and provision of human biospecimens used in this research. The authors thank the study participants of the RV217 Cohort.

## Disclaimer

The views expressed are those of the authors and should not be construed to represent the positions of the U.S. Army, the Department of Defense, the National Institutes of Health, the Department of Health and Human Services, or the Henry M. Jackson Foundation for the Advancement of Military Medicine, Inc. The investigators have adhered to the policies for protection of human subjects as prescribed in AR-70-25.

## Methods

### Human blood and lymph node samples

Whole blood from healthy blood donors was obtained from the New York Blood Center. Peripheral blood mononuclear cells (PBMCs) were isolated by Ficoll density gradient centrifugation using Ficoll-Paque™ PREMIUM (Cytiva, 45-001-751), cryopreserved in heat-inactivated fetal bovine serum (FBS; Gibco, A5670701) containing 5% dimethyl sulfoxide (DMSO; Sigma-Aldrich, D8418) and stored in liquid nitrogen until use.

Human lymph nodes were obtained from adults undergoing surgical resection at Tampa General Hospital (Tampa, Florida). All samples were collected in accordance with the Institutional Review Boards of the University of South Florida and Tampa General Hospital, and all participants provided written informed consent. Immediately after excision, lymph node tissue was placed in sterile phosphate-buffered saline (PBS; Gibco, 10010-023) supplemented with MycoZap Plus-PR (Lonza, VZA-2021) and transported to the laboratory on ice. Tissues were processed into single-cell suspensions within two hours of collection. After manual removal of surrounding fat, the lymph node was minced and digested in 40 mL of complete culture medium (RPMI 1640 GlutaMAX supplemented with 10% FBS and 1% penicillin/streptomycin) containing collagenase type IV (Gibco, 17104019; 1 mg/mL) and DNase I (Worthington Biochemical Corporation, NC9709009; 100 µg/mL) and incubated for 30 minutes at 37°C in a shaking incubator, followed by mechanical dissociation and passage through a 100 µm cell strainer (Corning, 431752). Recovered cells were cryopreserved in heat-inactivated FBS containing 5% DMSO and stored in liquid nitrogen until use. For culture, PBMCs and lymph node single-cell suspensions were thawed and resuspended in complete medium: RPMI 1640 GlutaMAX (Gibco, 72400047) supplemented with 10% FBS and 1% penicillin/streptomycin (Gibco, 15140122). All PBMC and lymph node donors were HIV-1 negative.

### HIV-1 infected study participants

PBMC samples, as well as CD4 and CD8 T cell count data of HIV-1 infected participants were from the RV217 Thailand cohort as we previously reported^25,26^. We analyzed the CD4 and CD8 count data from a total of 21 RV217 Thailand participants for whom we have previously determined coreceptor usage at around two years after HIV-1 transmission^25^. The CD4 and CD8 count data were from the same time points when coreceptor usage was determined. All participants were antiretroviral therapy (ART) naive at the time points analyzed.

### Cell lines

Human embryonic kidney (HEK) 293T cells were obtained from the American Type Culture Collection (ATCC CRL-3216) and TZM-bl cells were obtained from the NIH HIV Reagent Program (BEI Resources, HRP-8129). Both cell lines were maintained in Dulbecco’s modified Eagle medium (DMEM; Gibco, 11-965-092) supplemented with 10% FBS and 1% penicillin/streptomycin at 37 °C, 90% humidity and 5% CO_2_, and were split 1:10 with 0.05% trypsin/EDTA (Trypsin-EDTA; Gibco, 25300-054) when 70–80% confluent.

### HIV-1 infectious molecular clones

Full-length HIV-1 infectious molecular clones (IMCs) of two X4-tropic strains NL4.3 and SG3 and three R5-tropic strains CH058, CH040 and TRJO were obtained from the NIH HIV Reagent Program (BEI Resources, ARP-114, ARP-2003, HRP-11856, HRP-11740, HRP-11747). The full-length IMC of the X4 tropic strain 40700 was generated and characterized in our previous study^2^. Among the six HIV-1 strains, NL4.3 and SG3 are derived from chronic HIV-1 infection, while 40700, CH058, CH040 and TRJO are transmitted/founder (T/F) HIV-1 strains. All viruses are subtype B except for 40700 which is CRF01_AE.

### HIV-1 viral stock preparation and titration

HIV-1 viral stock was prepared and titrated as previously described^2,21^. In brief, 6 μg of IMC of each virus was transfected into 293T cells in a T25 flask using the FuGENE6 transfection reagent (Promega, E2691). The cells were cultured at 37°C for 6 h before the medium was completely replaced by fresh medium. The culture supernatants were harvested at 72 h post transfection, aliquoted and stored at -80°C until use. The infectious titer (TCID_50_) of the viral stock was determined in TZM-bl cells. The p24 concentrations of viral stocks were determined using the p24 ELISA kit (Revvity, NEK050B001KT).

### Microfluidic chip activation

Microfluidic chips (Chip-S1) and the culture module (ZOE-CM2) were obtained from Emulate Inc (Boston, MA USA). The Chip-S1 is a stretchable polydimethylsiloxane (PDMS) device consisting of two parallel microfluidic channels separated by a porous membrane (7 µm pore diameter): a top channel measuring 1 mm (width) × 1 mm (height) and a bottom channel measuring 1 mm × 0.2 mm. Before chip seeding, the inner surfaces of both channels were chemically activated to enable extracellular matrix (ECM) attachment, following the manufacturer’s protocol. Briefly, ER-1 working solution (0.5 mg/mL in ER-2 buffer; Emulate) was added to each channel and chips were exposed to UV light for 15 min. Chip channels were then aspirated and washed with ER-2 solution followed by PBS (Gibco, 10010-023). Chip channels were coated with an ECM solution containing 100 µg/mL Matrigel (Corning, 354234) and 100 µg/mL type I collagen (Corning, 354249) in PBS. The coated chips were incubated overnight at 4°C. The following day, the chips were transferred to a 37°C incubator for 2 h. Subsequently, the channels were vacuum-aspirated and washed twice with PBS, followed by vacuum aspiration to remove residual liquid. The chips were then incubated for 2 h at room temperature in a biosafety cabinet, followed by a 2 h incubation at 37°C. The chips were then ready for cell seeding.

### Lymphoid organ-chip culture

Cryopreserved PBMCs were thawed and counted on an Invitrogen automated cell counter (Countess 3) before chip seeding. The cells were subsequently resuspended on ice in an ECM gel containing 60% Matrigel (Corning, 354234), 15% type I collagen (Corning, 354249) and 25% complete RPMI 1640 GlutaMAX medium. Approximately 16 µL of ECM gel containing 8-10 × 10^6^ PBMCs was introduced through the inlet port of the bottom channel of a S1 chip, while degassed complete RPMI 1640 GlutaMAX medium was simultaneously flushed through the top channel inlet. Chips were incubated for 30 min at 37°C to allow gelation. Following gelation, both the inlet and outlet ports of the bottom channel were sealed with sterile polyethylene terephthalate (PET) discs to prevent fluid leakage and bubble formation in the bottom channel.

Before chip insertion, the inlet and outlet reservoirs of the top channel of each Pod (Emulate) were filled with 3 mL and 300 uL of fresh degassed complete RPMI medium, respectively, whereas the bottom channel reservoirs were left empty. The pods were primed in the ZOE-CM2 module using the prime cycle. Chips were then inserted into Pods, connected to the ZOE module, and perfused through the top channel inlet at 40 µL/h. The culture medium was recirculated daily by transferring effluent from the outlet reservoir to the inlet reservoir. Every 6 days, 50% of the perfused culture medium was replaced with a 1:1 mixture of retained conditioned medium and fresh complete culture medium.

### Lymphoid organ-chip infection and HIV-1 replication kinetics determination

Four days after cell seeding, chips were infected overnight with individual X4 tropic (40700, NL4.3, SG3) or R5 tropic (CH058, CH040, TRJO) HIV-1 strains at 5 ng p24 per 10^6^ cells (approximately 0.001 M.O.I.). During overnight infection, the flow rate was set at 20 µL/h using the ZOE-CM2 module. Following overnight infection, the top channel reservoirs were aspirated and washed four times with fresh culture medium. Each wash was performed over 2 h at an increased flow rate of 70 µL/h. After the fourth wash, the chips underwent an additional overnight wash at a flow rate of 20 µL/h to completely remove the residual virus in the inoculum stocks. The following day, the culture medium was completely aspirated from both the inlet and outlet reservoirs of the top channel and replaced with fresh pre-warmed culture medium. Effluent samples were collected on days 2, 5, 8 and 11 after infection to monitor HIV-1 replication. HIV-1 replication kinetics were assessed by measuring the p24 concentration in the effluent using HIV-1 p24 ELISA kit (Revvity, NEK050B001KT).

### Static 3D culture and HIV-1 infection

To compare cell phenotype and HIV-1 replication kinetics between microfluidic chip and conventional static 3D culture, 4 × 10^6^ PBMCs were seeded in the same ECM gel containing 60% Matrigel (Corning, 354234), 15% type I collagen (Corning, 354249) and 25% complete RPMI 1640 GlutaMAX medium at the same cell concentration as in lymphoid organ-chip (approximately 8 µL of ECM gel containing 4 × 10^6^ PBMCs). The cell-ECM suspension was cultured in 1 mL complete medium in a 24-well plate.

For HIV-1 infection, PBMCs recovered from static 3D culture were infected overnight in a U-bottom plate with the same dose of each HIV-1 strain as for the lymphoid organ-chip (5 ng p24 per 10^6^ PBMCs). Following overnight infection, cells were washed 3 times in culture medium and resuspended in the same ECM gel at the same cell concentration as used for chip seeding (approximately 8 µL of ECM gel containing 4 × 10^6^ PBMCs). The infected cell-ECM suspension was cultured in 1 mL complete medium in a 24-well plate. Culture supernatants were collected on day 2, 5, 8 and 11 after infection and HIV-1 replication kinetics were assessed by measuring the p24 concentration in culture supernatants using HIV-1 p24 ELISA kit (Revvity, NEK050B001KT).

### Cell recovery from chips

To recover cells from the chips, chips were disconnected from Pods, and the cells were recovered from both the bottom and top channels, since cells migrated from the bottom to the top channel during culture. The outlet ports of both channels were plugged with sterile empty tips, and 200 µL of Cell Recovery Solution (Corning, 354253) was manually perfused through the inlet ports of both channels. The recovered cells were collected and centrifuged at 500g for 5 min, resuspended in 1 mL Cell Recovery Solution and incubated for 1 hour at 4°C. Cells were then centrifuged again at 500g for 5 min, resuspended in PBS and counted on an Invitrogen automated cell counter (Countess 3) with viability dye before downstream analysis.

### Flow cytometry

For cell surface staining, cells were stained with a panel of fluorophore-conjugated antibodies (Fluorophore conjugates, catalog numbers and dilution factor of each antibody are listed in Supplementary Table 1): CD3 (OKT3), CD4 (SK3), CD8 (SK1), CD45RO (UCHL1), HLA-DR (L243), PD-1 (EH12.1), CCR5 (2D7), CCR7 (2-L1-A), CXCR4 (12G5), CXCR5 (RF8B2) and CD38 (HIT2). Each antibody was diluted 1:20 in FACS buffer for staining (PBS supplemented with 0.5% FBS and 2 mM EDTA (Invitrogen, AM9260G); 1 µl antibody in 19 µl FACS buffer) except for the CXCR4 antibody which was diluted 1:100 according to the results of antibody titration as we previously described^25^. Cells were stained for 30 min at room temperature in the dark. Following incubation, cells were washed once with FACS buffer and stained with the BD Horizon Fixable Viability Stain 660 (FVS660; BD Biosciences, 564405) for 15 min at room temperature in the dark. Cells were subsequently washed twice with the FACS buffer.

For Ki67 intracellular staining, cells were fixed using the BD Pharmingen^TM^ Transcription Factor Buffer Set (BD Biosciences, 562574). Briefly, cells were incubated in 1x TF Fix/Perm buffer for 40 min at 4°C, followed by two washes with the TF Perm/Wash buffer (BD Biosciences, 51-9008102). Intracellular staining was performed by incubating the cells with 5 µl of anti-Ki67 antibody (B56; BD Biosciences, 573363) diluted in 95 µl of TF Perm/Wash buffer (1:20 dilution) for 40 min at 4°C in the dark. Cells were subsequently washed twice with the TF Perm/Wash buffer and resuspended in FACS buffer before acquisition.

Samples were acquired on a NovoCyte Advanteon cytometer (Agilent) and analyzed using the FlowJo software (BD Biosciences). T cell subsets were defined as naïve (CCR7^+^CD45RO^−^), central memory (CCR7^+^CD45RO^+^) and effector memory (CCR7^−^CD45RO^+^); T follicular helper (Tfh) cells were defined as CXCR5^+^PD-1^+^ CD4 T cells and T follicular cytotoxic (Tfc) cells as CXCR5^+^PD-1^+^ CD8 T cells.

### Immunofluorescence microscopy

Immunofluorescence microscopy of the lymphoid organ-chip was performed as previously described^17^. In brief, four days after chip seeding, medium in the top channel was aspirated and chips were fixed by perfusing the top channel with 4% paraformaldehyde (Thermo Scientific, 043368.9M) in PBS and incubating for 1 h at room temperature on a rocker. Chips were then washed three times in PBS and incubated overnight at 4°C with primary antibodies against CD19-AF561 (eBioscience, 505-0199-42, HIB19, 1:25), CD4-AF488 (BD Biosciences, 557695, RPA-T4, 1:25) and ICOS (Cell Signaling Technology, 89601, D1KT2, 1:100) diluted in PBS containing 1% FBS, 0.1% Triton X-100 (Thermo Scientific, A16046.AE) and Fc block (Miltenyi Biotec, 130-059-901, 1:20). Following primary antibodies incubation, the chips were washed three times in PBS containing 2% FBS and 0.2% Triton X-100 on a rocker and incubated overnight at 4°C with an anti-rabbit IgG-AF647 secondary antibody (Invitrogen, A32733, polyclonal, 1:200). The chips were subsequently washed three times with the same washing buffer and stored in PBS at 4°C until imaging. Fluorescence imaging was performed on a BZ-X1000 fluorescence microscope (Keyence) using a 20x objective. The z-stack images were processed and analyzed by BZ-X Analyzer software (Keyence).

### Antiretroviral treatment (ART) in lymphoid organ-chip

Raltegravir, darunavir and nevirapine obtained from the NIH HIV Reagent Program (BEI Resources, HRP-11680, ARP-11447, and HRP-4666) were reconstituted in DMSO to a stock concentration of 10 mM. HIV-1 infection in lymphoid organ-chip was performed as described above, except that HIV-1 infection was performed 3 days after cell seeding (we compared viral replication kinetics when infection was performed three or four days after cell seeding and did not find significant difference). On day 5 post HIV-1 infection, ART was initiated in the designated chips using a drug cocktail containing raltegravir (10 µM), darunavir (10 µM) and nevirapine (10 µM). Drug stocks were diluted in complete RPMI medium immediately before use and perfused through the top channel inlet reservoir. Effluent was collected daily and completely replaced with fresh drug-containing medium to make sure that the effluent collected at the next time point contain only new virions released from the cells. HIV-1 infected chips without ART were used as controls. Treatment was maintained for 6 days before cells were recovered from the chips. HIV-1 replication kinetics in treated and untreated chips were determined by quantifying the HIV-1 RNA copy numbers in the effluent (as described below). At the end of treatment (day 6 after ART initiation), cells were recovered from all treated and untreated chips for cell phenotype analysis and integrated HIV-1 DNA quantification.

### Quantification of HIV-1 RNA viral load in chip effluent

To quantify HIV-1 RNA viral load in chip effluent, RNA was extracted from 100 µL chip effluent using Quick-RNA^™^ Viral Kit (Zymo Research, R1034). A total of 8.5 μL of extracted RNA was used for one-step RT-PCR using the Superscript III one-step RT-PCR system (Invitrogen, 12574026). The one-step RT-PCR was performed using the forward primer Pol F1 5′-TACAGTGCAGGGGAAAGAATA-3’ (nt 4809–4829 in HXB2) and the reverse primer Pol R1 5′-CTTCTTGGCACTACTTTTATGTCAC-3’ (nt 4993–5017) with the following PCR condition: a reverse transcription step at 55^◦^C for 30 min; A denaturing step at 94^◦^C for 2 min; 16 cycles of a denaturing step at 94 ^◦^C for 15 s, an annealing step at 55^◦^C for 30 s, an extension step at 68^◦^C for 1 min, and one cycle of an additional extension at 68^◦^C for 5 min. The first round PCR products were diluted 10-fold and a total of 6.4 μL of diluted PCR products were used for the real-time PCR using the forward primer Pol F1, the reverse primer Pol R2 5′-CTGCCCCTTCACCTTTCC-3’ (nt 4957–4974), and the probe Pol FamZen: 5’-/56-FAM/TTTCGGGTT/ZEN/TATTACAGGGACAGCAG/3IABk FQ/-3’ (nt 4896–4921). The real-time PCR was performed on the QuantStudio 3 Real-Time PCR Systems (Thermo Fisher Scientific) using the following conditions: A denaturing step at 95^◦^C for 4 min, 45 cycles of a denaturing step at 95^◦^C for 3 s, an annealing and extension step at 60^◦^C for 11 s. The copy number of the HIV-1 RNA was determined by using the RNA standard generated by *in vitro* transcription as described in our previous study^2^.

### Quantification of integrated HIV-1 DNA

Integrated HIV-1 DNA were quantified as previously described^2,51^. In brief, the absolute cell number was determined by amplifying the human CD3 gene using the primers HCD3OUT5 5′-ACTGACATGGAACAGGGGAAG-3′ and HCD3OUT3 5′-CCAGCTCTGAAGTAGGGAACATAT-3’. To quantify integrated HIV-1 DNA, the first round PCR was performed using the primers ULF1 5’-ATGCCACGTAAGCGAAACTCTGGGTCTCTCTDGTTAGAC-3’, Alu1 5′-TCCCAGCTACTGGGGAGGCTGAGG-3’ (nt 8674–8697), Alu2 5′-GCCTCCCAAAGTGCTGGGATTACAG-3’ (nt 6237–6261), HCD3OUT5 and HCD3OUT3. The first-round PCR conditions were as follows: a denaturation step at 95^◦^C for 8 min; 12 cycles of a denaturing step at 95^◦^C for 1 min, an annealing step at 55^◦^C for 1 min, an extension step at 72^◦^C for 10 min, followed by an elongation step at 72^◦^C for 15 min. The first round PCR products were diluted 10-fold and a total of 6.4 μL of the diluted PCR products were used for real-time PCR on the QuantStudio 3 Real-Time PCR Systems (Thermo Fisher Scientific) using TaqMan Fast Advanced Master Mix (Applied Biosystems, 4444557). The copy number of integrated HIV-1 DNA were quantified using the primers Lambda T 5′-ATGCCACGTAAGCGAAACT-3’ (nt 1555–1572), UR2 5′-CTGAGGGATCTCTAGTTACC-3’ (nt 583–602, 9668–9687), and the probe UHIV FamZen 5’-/56-FAM/CACTCAAGG/ZEN/CAAGCTTTATTGAGGC/3IABkFQ/-3’ (nt 522–546, 9607–9631). The copy numbers of human CD3 gene were determined using the primers HCD3IN5′ 5′-GGCTATCATTCTTCTTCAAGGT-3′, HCD3IN3′ 5′-CCTCTCTTCAGCCATTTAAGTA-3′, and the probe CD3 FamZen 5’-/56-FAM/AGCAGAGAA/ZEN/CAGTTAAGAGCCTCCAT/3IABkFQ/-3’. The real-time PCR conditions were as follows: A denaturing step at 95^◦^C for 4 min, 40 cycles of a denaturing step at 95^◦^C for 3 s, an annealing and extension step at 60^◦^C for 11 s. The ACH2 cell lysates were used for standard curve as previously described^2^.

### Single-genome amplification

Single-genome amplification (SGA) of 3’ half HIV-1 genome was performed as previously described^52^. In brief, viral RNA was extracted from chip effluent collected at day 14 after infection with CH058 using the Quick-RNA^™^ Viral Kit (Zymo Research, R1034). cDNA was synthesized with SuperScript^™^ III Reverse Transcriptase (Invitrogen, 18080-051) using primer 1.R3.B3R 5’-ACTACTTGAAGCACTCAAGGCAAGCTTTATTG-3’ (nt 9611-9642). The first-round PCR was performed using the primers 07For7 5’-CAAATTAYAAAAATTCAAAATTTTCGGGTTTATTACAG-3’ (nt 4875–4912) and 2.R3.B6R 5’-TGAAGCACTCAAGGCAAGCTTTATTGAGGC-3’ (nt 9636–9607), and the second-round PCR was performed with the primers VIF1 5’-GGGTTTATTACAGGGACAGCAGAG-3’ (nt 4900–4923) and Low2c 5’-TGAGGCTTAAGCAGTGGGTTCC-3’ (nt 9591–9612). Two microliters of the first round PCR products were used for the second round PCR amplification. The PCR conditions were as follows: one cycle at 94°C for 2 min; 36 cycles of a denaturing step at 94°C for 15 s, an annealing step at 60°C for 30 s, an extension step at 68°C for 4 min; and one cycle of an additional extension at 68°C for 10 min. The PCR amplicons were directly sequenced by the cycle sequencing and dye terminator methods. Individual sequences were assembled and edited using Sequencher (Gene Codes). All PCR amplifications were performed using Platinum^TM^ Taq DNA Polymerase High Fidelity (Thermo Fisher Scientific, 11304029).

### HIV-1 genetic and evolutionary analysis

HIV-1 envelope sequences were aligned by the Gene Cutter tool in the Los Alamos HIV Sequence Database (https://www.hiv.lanl.gov/content/sequence/GENE_CUTTER/cutter.html), followed by manual adjustment. Phylogenetic tree was constructed by MEGA6 using the maximum-likelihood method. Highlighter plot was generated using the Highlighter tool in Los Alamos HIV Sequence Database (https://www.hiv.lanl.gov/content/sequence/HIGHLIGHT/highlighter_top.html). HIV-1 evolution was analyzed using the Poisson Fitter tool in Los Alamos HIV Sequence Database (https://www.hiv.lanl.gov/content/sequence/POISSON_FITTER/pfitter.html).

## Statistical analysis

Experiments were performed using PBMCs from three different donors unless otherwise stated (for example, the ART experiment was performed in a single donor due to PBMC availability). Statistical analyses were performed using GraphPad Prims 11. Statistical comparisons were performed using unpaired two-tailed *t*-test or paired two-tailed *t*-test or two-tailed Mann Whitney test as described in figure legends. Differences between groups were considered statistically significant when *P* < 0.05.

